# Performance comparison: exome-sequencing as a single test replacing Sanger-sequencing

**DOI:** 10.1101/2020.11.29.400853

**Authors:** Hila Fridman, Concetta Bormans, Moshe Einhorn, Daniel Au, Arjan Bormans, Yuval Porat, Luisa Fernanda Sanchez, Brent Manning, Ephrat Levy-Lahad, Doron M. Behar

**Author notes:** **Corresponding author:**, Medical Genetics Institute, Shaare Zedek Medical Center, Jerusalem 91031, Israel, P. (972) 25645213, F. (972) 26666935.

## Abstract

Systematic performance comparing the results of exome-sequencing as a single test replacing Sanger-sequencing of targeted gene(s) is still lacking. In this study we compared Sanger-sequencing results of 258 genes to those obtained from next generation sequencing (NGS) using two exome-sequencing enrichment kits: Agilent-SureSelectQXT and Illumina-Nextera. Sequencing was performed on leukocytes and buccal-derived DNA from a single individual, and all 258 genes were sequenced a total of 11 times (using different sequencing methods and DNA sources). Sanger-sequencing was completed for all exons, including flanking ±8bp regions. For the 258 genes, NGS mean coverage was >20x for >98% and >91% of the regions targeted by SureSelect and Nextera, respectively. Overall, 449 variants were identified in at least one experiment, and 407/449 (90.6%) were detected by all. Of the 42 discordant variants, 23 were determined as true calls, summing-up to a truth set of 430 variants. Sensitivity of true-variant detection was 99% for Sanger-sequencing and 97%-100% for the NGS experiments. Mean false-positive rates were 3.7E-6 for Sanger-sequencing, 2.5E-6 for SureSelect-NGS and 5.2E-6 for Nextera-NGS. Our findings suggest a high overall concordance between Sanger-sequencing and NGS. Both methods demonstrated false positive and false negative calls and similar performances. Consequently, high clinical suspicion for a specific diagnosis should override negative results of either Sanger-sequencing or NGS.

## INTRODUCTION

Next Generation Sequencing (NGS) has taken the fast lane from the bench to the bedside. What was once a “last resort” technique for exceptional cases in which a molecular diagnosis could not be achieved by standards means has become a routine test allowing physicians to translate genomic information into clinically actionable decisions (1). NGS has been applied for a variety of purposes, including exome/genome-sequencing for diagnosis of genetic diseases (2, 3), cancer genomics (4), discovery of transcription factor binding sites (5), transcriptome profiling (6), DNA methylation sequencing (7) and noncoding RNA expression profiling (8). In the clinical context, the wide scope of testing enabled by NGS has facilitated the deciphering of the genetic bases of various human diseases for which the molecular mechanism was unknown (9), has expanded the number of genes known to be associated with defined phenotypes (10) and has led to detection of novel mutations in genes with well described functions (11, 12).

Molecular investigation of genetically heterogeneous phenotypes often mandates the sequential or simultaneous sequencing of multiple genes which can be achieved through Sanger-sequencing or NGS. Sanger-sequencing entails the design of specific primers to guarantee successful amplification of regions of interest and may overcome difficulties such as sequencing within GC-rich regions and separating sequencing of genes from their pseudogenes (13, 14). However, Sanger-sequencing of multiple genes for diagnostic purposes is costly and time consuming, whereas it is eminently feasible using NGS (15). The advantages of NGS platforms over Sanger-sequencing with respect to genotyping capacity and cost have been demonstrated repeatedly (1). Indeed, NGS applications are readily available as gene panels or through exome-sequencing (16) for analysis of various phenotypes, e.g. non-syndromic deafness, neuro-muscular diseases and Noonan-spectrum syndromes (17–20). However, NGS also has shortcomings. NGS solutions have been previously associated with higher error rates as compared to Sanger-sequencing (21), which led to the common practice of confirming NGS results by Sanger-sequencing. Other common problems include unequal coverage throughout the targeted region, sequencing of pseudogenes, and difficulty in sequencing GC-rich regions (22). It is therefore surprising that unbiased, two-way systematic comparisons of performance between various NGS platforms and Sanger-sequencing are still rare, despite the important implications of such comparisons for genetic diagnostics.

Previous studies examined the ability of NGS to detect variants identified by Sanger-sequencing. For a panel comprised of a limited number of genes for hereditary colon cancer, arrhythmias, cardiomyopathies, and other cardiovascular-related genes, it was shown that all 919 variants previously observed by Sanger-sequencing were also identified by NGS (23). Another study, however, demonstrated that up to 18% of 137 pathogenic variants identified by Sanger-sequencing, in the context of neuromuscular diseases, were not identified by exome-sequencing (17). In another study, variants identified by clinical Sanger-sequencing were compared to results obtained from exome-sequencing for the same patients (24). exome-sequencing identified 97.3% of the coding variants and 81.8% of the non-coding variants detected by Sanger-sequencing. Nine genes were excluded from the comparison due to consistently low coverage in exome-sequencing. These results demonstrated molecular circumstances in which Sanger-sequencing performed better than NGS (24). Neither study examined performance of NGS independently of the Sanger-sequencing results, e.g. if there were additional variants identified by NGS that were missed by Sanger-sequencing. In fact, all these studies were one-way comparisons that assessed sensitivity of NGS for variants identified by Sanger-sequencing. None provided data regarding NGS false positives or Sanger-sequencing false negatives.

We present a comparative, unbiased evaluation of sequencing results of 258 genes comprising ~1.3% of the human exome. DNA from a single individual was extracted from peripheral blood leukocytes and buccal-swabs. The platforms assessed were Sanger-sequencing of the coding exons ±8bp of the 258 genes, the Agilent SureSelect^QXT^ exome capture kit (SureSelect) and the Illumina Nextera Rapid Capture Expanded Exome (Nextera). We aimed to shed light on the performance and accuracy of each of the sequencing methods, from the two DNA sources, and to assess the implications of replacing Sanger-sequencing with exome-sequencing, using standard capture kits. Although only one individual was sequenced, to ensure a single reference sequence, we performed 11 different sequencing experiments for each of 258 genes, using different sequencing strategies and different DNA sources. This enabled in-depth comparisons between Sanger-sequencing and NGS results.

## MATERIALS AND METHODS

### Samples and sequencing

Samples from blood and buccal-cells were taken the same day from one healthy individual following informed consent. All DNA handling and sequencing procedures were completed at Gene by Gene’s CAP accredited laboratory in Houston, Texas. DNA was extracted from blood (peripheral leukocytes) and buccal-swabs following standard protocols, and quantitated with a SpectraMax190 (Molecular Devices, Sunnyvale, CA). Sanger-sequencing was completed for 258 genes on DNA extracted from buccal-cells by amplifying the coding exons and their flanking regions (approximately 20bp from each side) using conventional PCR techniques. PCR products were purified using magnetic-particle technology (Seradyn, Inc.). After purification, all fragments were sequenced with forward and reverse primers. Sequencing was performed on a 3730xl DNA Analyzer (Applied Biosystems), and the resulting sequences were analyzed with the Sequencher software (Gene Codes Corporation). Variants were analyzed relative to the reference sequences deposited in the National Center for Biotechnology Information.

exome-sequencing was performed using the Agilent enrichment capture kit (SureSelect^QXT^), and the Nextera enrichment rapid capture kit (FC-140-1006) on DNA extracted from both peripheral leukocytes and buccal-cells. exome-sequencing data were generated using the Illumina HiSeq2500 with manufactures’ protocols for all runs. A coverage of 70x was considered the minimal threshold for clinical grade sequencing.

### Sequencing quality reassurance

For Sanger-sequencing, each nucleotide was covered by at least two different sequences, preferably by one forward and one reverse sequences. In cases where this was not possible due to poly-regions, two sequences from the same direction were generated independently. The interpretation of the data was done by two operators independently. We used a Phred score of 30 as our confidence threshold.

### Exome-sequencing Data Analysis and Quality Reassurance

Each exome-sequencing sample was analyzed using the Genoox platform (http://www.genoox.com). The NGS pipeline was based on BWA aligner (25), and the two variant callers GATK HaplotypeCaller (26) and FreeBayes (27). Coverage reports were obtained from Genoox platform. The minimal Mapping Quality (MQ) was MQ>0 for each read. The main parameters used for filtering were identical for both variant callers and comprised a quality score>100, a depth (DP)>4 and the quality/depth ratio (QD) of 7.

### Variants comparison

We included 258 genes in our analysis (*Table S1*). The list of genes was assembled from the genes previously Sanger-sequenced at Gene by Gene’s CAP accredited laboratory. The only genes excluded were those expected to perform less efficiently in NGS, due to issues involving pseudo genes or GC-rich regions (i.e*. GBA* (OMIM 606463)*, MAPT* (OMIM 157140)*, PMS2* (OMIM 600259)). The analyzed regions include coding regions (exons)±8bp flanking introns attempting to cover the splice sites. Following quality reassurance, we established the list of all the variants identified by any of the different platforms and experiments, and checked systematically how many experiments detected each variant. Variants that were detected by all experiments were classified as true variants. Variants that were not concordant between all experiments were considered potential false positives or false negatives. The Sanger-sequencing suspected false positive or negative variants were re-sequenced with an alternative primer pair. The NGS suspected false positive variants were checked for their quality parameters (DP<20; QD<7; QUALITY<100).

## RESULTS

We compared the results of Sanger-sequencing and exome-sequencing for 258 genes. We analyzed two exome-sequencing runs. In the first run, exome-sequencing was performed on DNA from both sources (leukocyte and buccal), once using the SureSelect kit and once using the Nextera kit. Each exome-sequencing was performed in duplicate, so this run included 8 exome-sequencing tests, enabling within-run comparisons. The second run included two exome-sequencing tests performed on leukocyte-extracted DNA, one using the SureSelect kit and one using the Nextera kit. This enabled between-run comparisons.

### Targeted genomic region

The targeted genomic region, as noted above, included 258 genes (*Table S1*), with a total of 4,629 exons. The genomic region encompassed by these exons is 729,724 bps; 803,788 bps including ±8bp of the intron/exon boundaries. *Table 1* summarizes the targeted genomic region, the expected overlap according to the SureSelect and Nextera BED files, and the actual adequately enriched regions (>20x) obtained from the experiments. More than 98%/91% of the region was covered with >20x coverage for the SureSelect/Nextera experiments.

**Table 1.** NGS results: expected vs. actual enriched genomic regions

| Enrichment capture-kit | SureSelect |  |  |  |  | Nextera |  |  |  |  |
| --- | --- | --- | --- | --- | --- | --- | --- | --- | --- | --- |
| Expected enriched region (exons)<sup>a</sup> | 98.4% |  |  |  |  | 99.0% |  |  |  |  |
| Expected enriched region (exons±8bp)<sup>a</sup> | 98.2% |  |  |  |  | 90.4% |  |  |  |  |
| Sequencing run | Run 1 |  |  |  | Run 2 | Run 1 |  |  |  | Run 2 |
| DNA source<sup>b</sup> | Buccal Dup A | Buccal Dup B | Blood Dup A | Blood Dup B | Blood | Buccal Dup A | Buccal Dup B | Blood Dup A | Blood Dup B | Blood |
| Actual enriched region >20X (exons)<sup>c</sup> | 98.9% | 99.2% | 98.8% | 98.7% | 98.4% | 91.9% | 92.1% | 95.2% | 94.3% | 95.0% |
| Actual enriched region >20X (exons±8bp)<sup>c</sup> | 98.9% | 99.1% | 98.8% | 98.6% | 98.4% | 91.3% | 91.5% | 94.9% | 93.9% | 94.7% |

<sup>a</sup> The percentage of nucleotides encompassing the 258 genes that are expected to be enriched, based on the respective BED file declared by the manufacturers.
<sup>b</sup> Dup=duplicate
<sup>c</sup> The percentage of nucleotides in the enriched region actually sequenced at >20x coverage

### Overall NGS performance

We performed ten NGS experiments. *Table 2* highlights the pivotal metrics including the mean coverage obtained for the exome-sequencing as well as the targeted region, and the percentage of bps obtained at a coverage of 0x, ≤10x, ≤20x and >20x.

**Table 2.** NGS coverage: comparison of capture-kits, DNA source and consistency.

| Enrichment capture-kit | SureSelect |  |  |  |  | Nextera |  |  |  |  |
| --- | --- | --- | --- | --- | --- | --- | --- | --- | --- | --- |
| Sequencing run | Run 1 |  |  |  | Run 2 | Run 1 |  |  |  | Run 2 |
| DNA source<sup>a</sup> | Buccal Dup A | Buccal Dup B | Blood Dup A | Blood Dup B | Blood | Buccal Dup A | Buccal Dup B | Blood Dup A | Blood Dup B | Blood |
|  | Mean coverage |  |  |  |  | Mean coverage |  |  |  |  |
| Whole exome | 174x | 193x | 201x | 163x | 135x | 90x | 97x | 106x | 96x | 75x |
| Enriched region<sup>b</sup> (exons) | 258x | 287x | 297x | 243x | 229x | 98x | 106x | 116x | 105x | 102x |
| Enriched region<sup>b</sup> (exons±8bp) | 254x | 282x | 292x | 238x | 225x | 97x | 104x | 114x | 103x | 101x |
|  | Coverage distribution <sup>c</sup> |  |  |  |  | Coverage distribution <sup>c</sup> |  |  |  |  |
| 0 | 0.05% | 0.07% | 0.14% | 0.13% | 0.09% | 0.57% | 0.62% | 0.56% | 0.70% | 0.64% |
| 0-10x | 0.46% | 0.33% | 0.57% | 0.52% | 0.59% | 2.61% | 2.49% | 1.93% | 2.10% | 1.69% |
| 10x - 20x | 0.58% | 0.48% | 0.52% | 0.70% | 0.91% | 5.53% | 5.38% | 2.65% | 3.30% | 2.94% |
| > 20x | 98.91% | 99.12% | 98.77% | 98.65% | 98.40% | 91.29% | 91.51% | 94.86% | 93.90% | 94.73% |

<sup>a</sup> Dup=duplicate
<sup>b</sup> The regions encompassing the targeted region comprising the 258 genes, based on the respective BED file as declared by the manufacturers.
<sup>c</sup> Percentage of bp in each coverage category, calculated for the exons±8bp of the enriched region.

Comparison of duplicate experiments within run 1, and comparison of run 1 and run 2 demonstrated no statistically significant differences for the major displayed metrics. For the entire exome, mean coverage was 166x with the SureSelect kit and 90x with the Nextera kit. In the targeted region (258 genes, including both the exons and the exon±8bp intron/exon boundaries), mean coverage of the SureSelect kit was about double that of the Nextera kit (*Table 2*). Notably, the total number of aligned reads was similar in both capture kits (>99%), but the SureSelect kit mean coverage was about double that of the Nextera kit mean coverage since a large proportion of Nextera reads did not align to the target region. For both the SureSelect and the Nextera capture kits, less than 1% of the nucleotides had 0x coverage in any of the experiments. The mean proportion of nucleotides with 0x coverage across all experiments of the same kit were 0.1% and 0.62% for the SureSelect and Nextera, respectively.

### Identified variants

We analyzed the exome-sequencing data using standard NGS analytic pipelines with predefined quality thresholds, as described in the *Methods* section. A total of 449 variants were identified at least once, in a total of 258 genes (range of 0-13 variants per gene) (*Table 3, Table S2, Table S3*). The majority of those variants, 407/449 (90.6%), were detected by all platforms and experiments. The remaining 42/449 (9.4%) discordant variants were examined further to determine whether they represent true or false variants (*Table 3, Table S3*).

**Table 3.** Discordant variants^a^

| Gene | cDNA | rs number | True variant<sup>b</sup> | Conclusion<sup>b</sup> |
| --- | --- | --- | --- | --- |
| *ABCC6* | c.841A>G | rs4780606 | N | Sanger FP |
| *ABCC6* | c.855C>T | rs4780605 | N | Sanger FP |
| *ABCC6* | c.955A>G | rs72657699 | N | Sanger FP |
| *IQCB1* | c.1301G>A | rs17849995 | Y | Sanger FN |
| *OPA1* | c.473G>A | rs7624750 | Y | Sanger FN |
| *TSC2* | c.5202T>C | rs1748 | Y | Sanger FN |
| *TSC2* | c.5397G>C | rs1051771 | Y | Sanger FN |
| *CFTR* | c.1251C>A | rs4727853 | N | NGS FP |
| *NOTCH3* | c.2742A>G | rs1043997 | N | NGS FP |
| *CACNA1A* | c.6669T>C | rs16051 | Y | Nextera FN |
| *DLL3* | c.515T>G | rs8107127 | Y | NGS FN |
| *DLL3* | c.546C>G | rs8106337 | Y | NGS FN |
| *PIK3R2* | c.700A>C | rs2241088 | Y | NGS FN |
| *COL5A1* | c.4560C>T | rs2228559 | Y | SureSelect bl-R1DA FN |
| *COL6A2* | c.1333-8T>C | rs73159701 | Y | SureSelect bu FN |
| *CDH1* | c.48+6C>T | rs3743674 | Y | Nextera FN |
| *FRMD7* | c.1101T>C | rs7051368 | Y | Nextera FN |
| *MEN1* | c.435C>T | rs61736636 | Y | Nextera bl FN |
| *FGF3* | c.69G>T | rs41538178 | Y | Nextera bl-R1DA FN |
| *GLIS3* | c.1270T>C | rs806052 | Y | Nextera bl-R1DB FN |
| *COL6A2* | c.2979C>T | rs6652 | Y | Nextera bl-R1DB FN |
| *LMX1B* | c.326+7G>C | rs1336980 | Y | Nextera R2 FN |
| *NF1* | c.702G>A | rs1801052 | Y | Nextera bu-R1DB FN |
| *COL5A2* | c.315C>A | rs4128539 | Y | Nextera bu-R1DB FN |
| *EIF2AK3* | c.407C>G | rs867529 | Y | Nextera bu-R1DA FN |
| *MESP2* | c.531G>A | rs75049807 | Y | Nextera bu-R1DB FN |
| *MYH7* | c.1095G>A | rs735711 | Y | Nextera bu-R1DA FN |
| *USH2A* | c.3812-8T>G | rs646094 | Y | Nextera bu-R1DA FN |
| *GPC3* | c.338-5delT | rs370737647 | N | SureSelect bu-R1DA FP |
| *MESP2* | c.558G>A | rs28546919 | N | SureSelect R2 FP |
| *CFTR* | c.1365G>T | rs79074685 | N | Nextera bl-R1DA FP |
| *COL1A1* | c.3046-6_3046-5delCT | rs138425306 | N | Nextera bl-R1DA FP |
| *PIGN* | c.2620-5delT | rs11437076 | N | Nextera bl-R1DA FP |
| *APP* | c.10G>T | ... | N | Nextera R2 FP |
| *FRMD7* | c.575A>C | rs746689690 | N | Nextera R2 FP |
| *NOTCH3* | c.4215C>T | ... | N | Nextera R2 FP |
| *NOTCH3* | c.4208G>T | ... | N | Nextera R2 FP |
| *SCNN1B* | c.275T>C | rs1596860414 | N | Nextera R2 FP |
| *RAG1* | c.461delT | ... | N | Nextera bu-R1DA FP |
| *FOXC1* | c.1359_1361delCGG | rs398123612 | N | Nextera bu-R1DB FP |
| *EFNB1* | c.*731_*732insC | rs61387446 | N | Nextera bu-R1DB FP |
| *EFNB1* | c.*732_*733insCAC | rs58007755 | N | Nextera bu-R1DB FP |

<sup>a</sup> Shaded: Variants discordant between the Sanger and NGS experiments (N=13).
Not shaded: Variants discordant within the various NGS experiments (N=29).
<sup>b</sup> bl-blood; bu-buccal; D-duplicate; FP-False positive; FN-False negative; R-run; N-no; Y-yes.

Discordant variants were classified into two groups: variants discordant between Sanger-sequencing and NGS (Sanger-NGS discordance) and variants discordant within NGS experiments (within-NGS-discordance). Sanger-NGS discordance was defined as discordance between Sanger-sequencing and most (≥5) NGS experiments. Within-NGS discordance was defined as discordance between NGS experiments, where the majority of NGS results (>5) were concordant with Sanger-sequencing results. Thirteen variants were Sanger-NGS discordant and 29 variants were within-NGS discordant (*Table 3, Table S3*). Among the 13 Sanger-NGS discordant variants, four variants (in the *DLL3* (OMIM 602768), *CACNA1A* (OMIM 601011) and *PIK3R2* (OMIM 603157) genes) represent NGS false negatives due to coverage failure (<10x), since they were identified by Sanger-sequencing and by all NGS experiments with sufficient coverage. In the other nine Sanger-NGS discordant variants, there was sufficient coverage in all NGS experiments. For these variants, we repeated Sanger-sequencing using an alternative primer pair.

Discrepancies were resolved for seven of the nine variants, showing that the original Sanger-sequencing included four false negative and three false positive calls. All three false positives were in the same PCR amplicon of the same gene (*ABCC6* (OMIM 603234)), and may be explained by individual variants in the primer region. The two remaining variants (*CFTR* (OMIM 602421) and *NOTCH3* (OMIM 600276) genes) were not detected by repeated Sanger-sequencing despite several primer redesign attempts, and were thus considered as NGS false positive calls. Twenty-nine variants were within-NGS discordant. Of these, 14 variants were not detected by the Sanger-sequencing and detected by only one of all NGS experiments: eight in leukocyte-derived DNA and four in buccal-swab DNA with Nextera capture; one in leukocyte DNA and one in buccal-swab DNA with SureSelect capture. These variants barely passed the quality filters, and had coverage<20 or Quality<100 or QD<7, and are therefore considered as NGS false positives due to low quality. Of the remaining 15 variants, 13 were detected by the Sanger-sequencing, the SureSelect experiments and at least one Nextera experiment. The other two variants were detected by the Sanger-sequencing, the Nextera experiments and at least one SureSelect experiment. All these 15 variants are thus considered as NGS false negatives. In 12 of them, lack of detection is explained by low coverage (<20x).

Cumulatively, we determined that of the 42 discordant variants, 23 were true variant calls, including four Sanger-sequencing false negatives and 19 NGS false negatives. The remaining 19 variants were false positives, including three Sanger-sequencing calls and 16 NGS experiments calls (*Table 3, Table S3*). False positive NGS calls included 9/16 with poor coverage (<20x), and 7/16 variants with >20x coverage, but borderline quality (QD<7). Notably, ten of the 14 NGS false positives (71.4%) have rs IDs. Thus, together with the 407 variants detected by all experiments, a total of 430/449 (95.8%) variants were defined as “true variants”, and regarded as the truth set for further analysis.

### Comparative Sequencing Performance

As shown in *Figure 1*, overall, sensitivity was >97% in all platforms and experiments. Sanger-sequencing had a detection rate of 99%. In NGS experiments, higher coverage resulted in better detection rates. The mean sensitivity of SureSelect and Nextera experiments was 99.5% and 98.5%, respectively.

**Figure 1.**
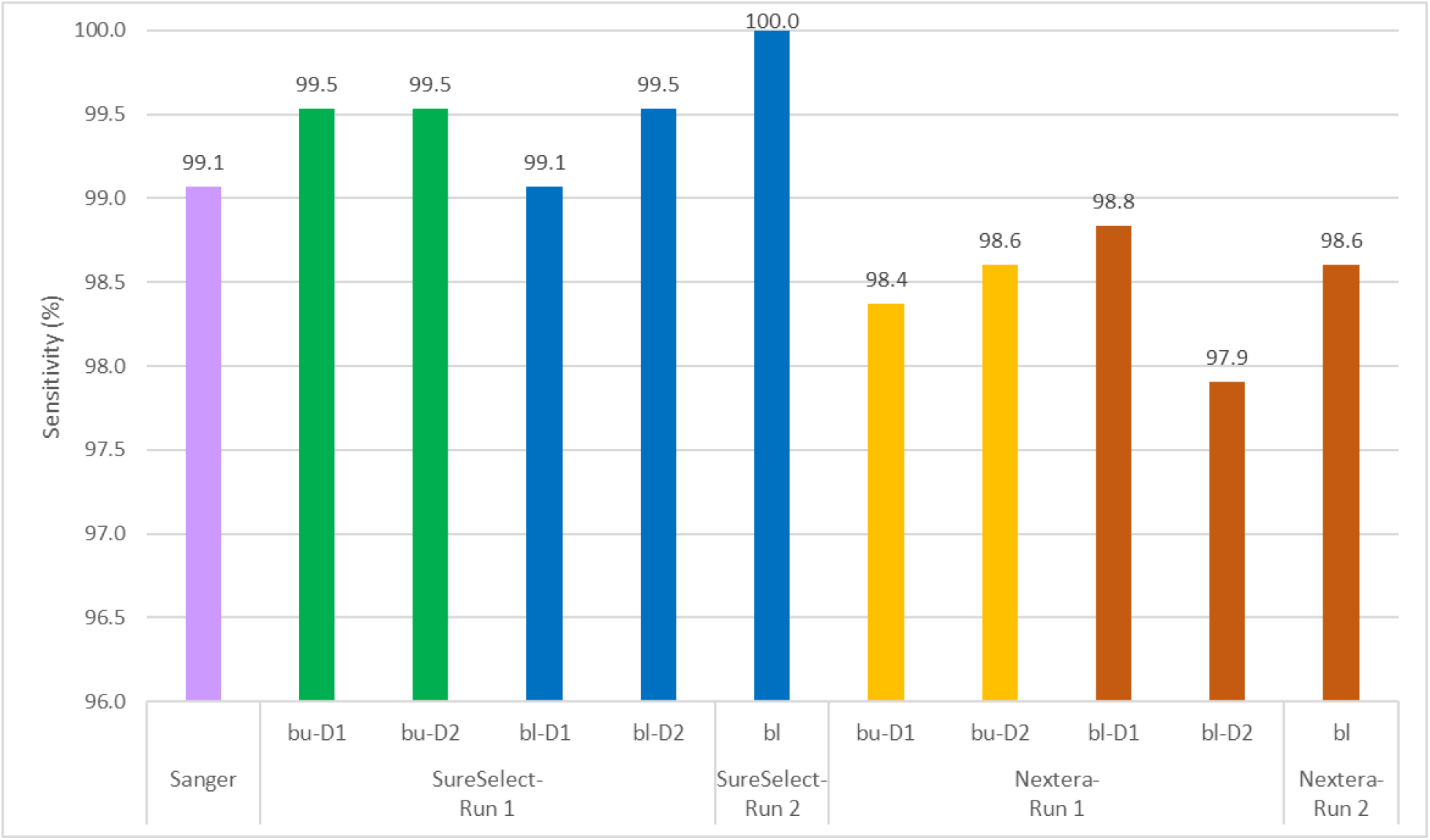
Sensitivity across all platforms and experiments. The percentage of 430 “true variants” detected in each experiment. bu-buccal; bl-blood; D-duplicate.

*Figure 2* presents the false positive and false negative calls for each of the platforms and experiments. Sanger-sequencing resulted in three false positives (all in the same PCR amplicon) and four false negative calls. The number of false negative calls observed in the Nextera experiments ranged between five and nine calls (mean of 6.6). Similarly, we detected 0-4 false negative calls in the SureSelect experiments (mean of 2). In general, the number of false negative calls decreased with higher coverage. None of the differences was statistically significant using Chi-square test (p>0.05). The Nextera experiments demonstrated 2-7 false positive calls (mean of 4.2), while the SureSelect experiments yielded 1-3 false positive calls (mean of 2). The PPV (positive predictive value) and NPV (negative predictive value) for all experiments were both >0.99.

**Figure 2.**
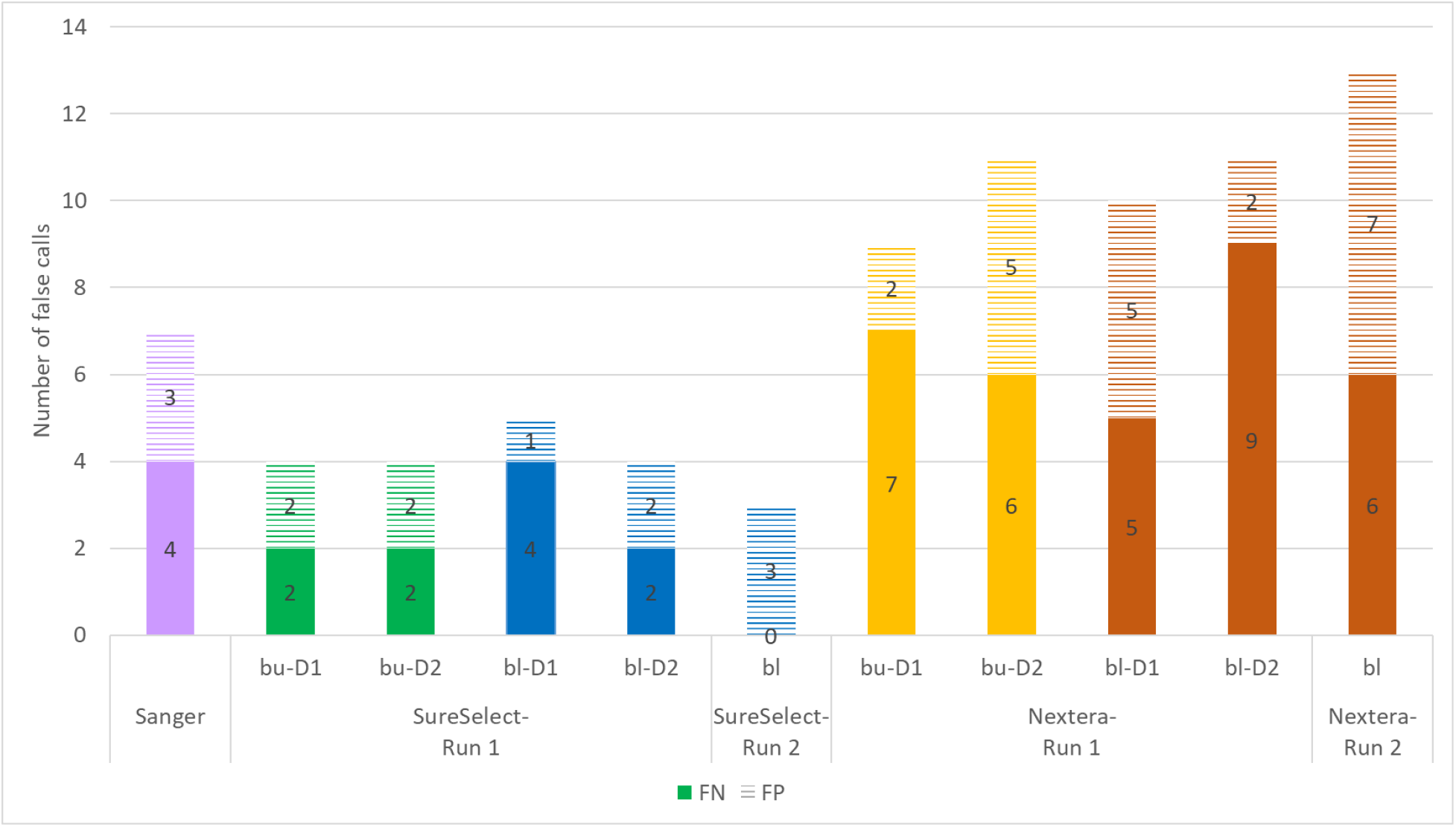
Number of false calls. The number of false positive and false negative variants is shown for each experiment. False negative calls-solid bars. False positive calls-striped bars. Number of each type of call is indicated within each bar. Bu-buccal; bl-blood; D-duplicate. None of the differences are statistically significant.

### Zygosity discordance between platforms and experiments

We observed zygosity discordance for a total of 10 variants (*Table 4*). In 5/10 the discordance was between Sanger-sequencing and all NGS experiments (Sanger-NGS discordance). In four of those five cases, Sanger-sequencing suggested a homozygous state while all NGS experiments suggested heterozygosity. In the fifth case, the opposite was noted. The NGS coverage for these variants ranged from 27x to 307x (average 183x). For the other 5/10 variants there was within-NGS zygosity discordance: Sanger-sequencing and SureSelect experiments were concordant but there was discordance with one of the Nextera experiments. Coverage was <20x in four of the cases, and 41x in the fifth case (*Table 4*).

**Table 4.** Zygosity discordance between Sanger and NGS^a^

| Gene | dbSNP | Sanger<br>zygosity | SureSelect<sup>b</sup> |  |  |  |  | Nextera<sup>b</sup> |  |  |  |  |
| --- | --- | --- | --- | --- | --- | --- | --- | --- | --- | --- | --- | --- |
|  |  |  | Run 1 |  |  |  | Run 2 | Run 1 |  |  |  | Run 2 |
|  |  |  | Buccal<br>Dup A | Buccal<br>Dup B | Blood<br>Dup A | Blood<br>Dup B | Blood | Buccal<br>Dup A | Buccal<br>Dup B | Blood<br>Dup A | Blood<br>Dup B | Blood |
| *ABCA3* | rs13332514 | hom | het<br>152/146 | het<br>147/157 | het<br>115/125 | het<br>111/116 | het<br>108/84 | het<br>121/112 | het<br>149/121 | het<br>153/100 | het<br>102/108 | het<br>69/77 |
| *ABCA4* | rs4847281 | het | hom<br>0/100 | hom<br>1/156 | hom<br>0/156 | hom<br>0/93 | hom<br>0/83 | hom<br>0/50 | hom<br>0/50 | hom<br>0/77 | hom<br>0/44 | hom<br>0/74 |
| *ATP13A2* | rs761421 | hom | het<br>34/24 | het<br>41/52 | het<br>22/31 | het<br>21/16 | het<br>10/17 | het<br>29/49 | het<br>37/45 | het<br>21/23 | het<br>33/27 | het<br>42/44 |
| *ATP13A2* | rs9435659 | hom | het<br>32/41 | het<br>57/72 | het<br>37/40 | het<br>27/32 | het<br>21/21 | het<br>124/163 | het<br>155/152 | het<br>106/130 | het<br>84/100 | het<br>71/84 |
| *B3GLCT* | rs1041073 | hom | het<br>86/76 | het<br>86/86 | het<br>92/93 | het<br>79/60 | het<br>91/58 | het<br>135/124 | het<br>128/143 | het<br>118/88 | het<br>113/76 | het<br>109/85 |
| *MESP2* | rs75049807 | het |  |  |  |  |  |  |  |  | hom<br>0/6 |  |
| *PIGN* | rs34227891 | hom |  |  |  |  |  | het<br>6/35 |  |  |  |  |
| *FGF3* | rs41538178 | het |  |  |  |  |  |  |  |  | hom<br>0/4 |  |
| *SCN1A* | rs7580482 | hom |  |  |  |  |  |  |  |  | het<br>1/10 |  |
| *SCNN1B* | rs238547 | hom |  |  |  |  |  |  |  |  | het<br>1/5 |  |

<sup>a</sup> hom=homozygote; het=heterozygote; dup=duplicate.
<sup>b</sup> For each variant, the numbers below the zygosity indicate the number of reference calls (left) and the number of variant calls (right). Results are shown only for NGS experiments discordant with Sanger sequencing.

We repeated Sanger-sequencing using an alternative primer pair for all 10 variants. The zygosity status changed for the first 5 variants, which indicates Sanger-sequencing error due to primer design in the first sequencing. The zygosity status of the other 5 variants did not change, which indicates a single NGS experiment error.

## DISCUSSION

Accuracy of NGS platforms and the ability to obtain full exome/genome-sequencing information are critical elements of clinical genetic testing and of precision medicine initiatives (28). We compared the performance of Sanger-sequencing and NGS for 258 genes that are commonly sequenced in a commercial laboratory. Sequencing these genes is generally requested as part of routine clinical testing for well-defined OMIM phenotypes or during the investigation of less defined, orphan phenotypes with unknown molecular bases. OMIM phenotypes are assigned to 257 of 258 genes included in our study (https://omim.org/) and 30 are among the 67 genes listed in the American College of Medical Genetics and Genomics recommendations for reporting of incidental findings in clinical exome and genome-sequencing (29). Accordingly, the genes we examined can be regarded as representative of common clinical scenarios mandating single or multiple gene sequencing. These scenarios often present the dilemma of choosing between Sanger-sequencing of specific gene(s), NGS of a gene-panel including the requested gene(s), or full exome-sequencing that might serve both immediate and future needs of the patient. Compared to exome-sequencing, sequencing specific gene(s) by either Sanger or NGS, involves higher sequencing costs per gene, and incurring additional costs of further genetic testing if a molecular diagnosis is not confirmed in the first round of testing. However, sequencing of single genes or gene panels are more likely to provide complete coverage of the targeted genes.

Although we only sequenced one individual in this study, the study we performed is unique in providing bidirectional comparison of Sanger-sequencing and NGS. By using different sequencing strategies, different NGS solutions and different DNA sources, each gene was sequenced a total of 11 times. This way we could determine with confidence whether each variant observed represents a true or false call, examine false negative and false positive rates of the methods, and evaluate characteristics of both true and false calls.

We first examined the effect of the DNA source and capture kit on NGS results, as well as consistency of NGS results between duplicates and between different runs. NGS experiments were performed using the exact guidelines of the respective manufacturers. Within each of the kits, it is evident that no statistically significant differences were noted between duplicates in the same run or different sequencing runs (*Table 2*). Differences in performance between capture kits have been previously reported (30–32). Our results demonstrate that the mean coverage obtained by the SureSelect kit was higher, reaching 182x/183x for DNA extracted from leukocytes or buccal-cells, respectively. There were no significant differences in the performance of NGS on DNA extracted from these two different sources.

We then determined the rates and types of false calls in both Sanger-sequencing and NGS. Of the 449 variants observed in the 258 genes analyzed, 90.6% (407) were detected in all sequencing experiments. Among the 42 variants which were not universally detected, we determined that 23 were true calls, defining a truth-set of 430 variants. The detection rate of true variants was 99.1% (426/430) for Sanger-sequencing, and ranged between 97.9% and 100% in the NGS experiments (*Figure 1*). Although the sensitivity, false positive and false negative rates calculated for all experiments were not statistically different, probably due to the small number of discordant variants, the differences observed offer important insights into the strengths and weaknesses of each sequencing method.

For Sanger-sequencing, the false positive rate was 3.7E-6, and the false negative rate was 0.009. Sanger false positives all occurred in one gene (*ABCC6* (OMIM 603234)), whereas false negatives occurred in three different genes. All false Sanger calls were resolved by using alternative primer pairs, suggesting they could be idiosyncratic to polymorphisms in the tested individual.

For NGS, the mean false positive rate was 2.5E-6 in the SureSelect experiments, and 5.2E-6 in the Nextera experiments. The mean false negative NGS rates were 0.005 for SureSelect experiments, and 0.01 for Nextera experiments. Most false positive NGS calls were singular events, i.e. they occurred in only one of 10 NGS experiments. These variants had barely passed the quality filter, and failed one or more of the basic quality parameters (DP>20; QUALITY>100; QD>7). Changing the quality filter thresholds might obviously decrease the number of false positives, but this would come at the cost of an increase in false negatives. Interestingly 15/19 (78.9%) of false positive calls, including all three Sanger-sequencing false positives, have rs IDs. This may indicate that current databases contain false positive calls presented as true variants. Conversely, all 430 true variants were detectable by NGS, as long as coverage was adequate and quality parameters were fulfilled, consistent with previous reports that there is no need for Sanger-sequencing confirmation of such high-coverage/high-quality variants (33).

NGS false negatives were all associated with low (<20x) coverage. In some cases, low coverage was a singular event (e.g. *COL5A* (OMIM 120215) or *COL5A2* (OMIM 120190)), but in other cases low coverage was systematic to a specific kit (e.g. *PIK3R2* (OMIM 603157) in SureSelect or *FRMD7* (OMIM 300628) in Nextera), or to both capture methods (e.g. *DLL3* (OMIM 602768)). Systematic false negatives indicate problematic genomic regions in which variant detection is hampered. This is a concern, since it was previously shown that large areas of medically actionable genes fall within low confidence regions (23), which might explain consistently low coverage for some variants. In this study, using exome-sequencing to target a specific set of genes encompassing ~1.3% of the exome, resulted inadequate coverage (>20x) of >98% and >91%, of exons±8bp with the SureSelect and the Nextera kits respectively (*Table 1*). This suggests that ~16,075 and ~72,340 bp (of a total of 803,788 bp in 258 genes), would not be adequately covered. Clearly, disease causing variants could be located in such regions with insufficient coverage. Taken together, our data suggest that exome-sequencing could miss ~2-3% of the coding variants and up to 7% of the non-coding variants. These results are in agreement with Hamilton et al (24) who demonstrated a 92.3% concordance between exome-sequencing and Sanger-sequencing in the ±20bp intronic/exon boundaries.

We also found zygosity status errors for both Sanger-sequencing and NGS. As presented in *Table 4*, Sanger-sequencing determined a false zygosity status of 5 variants (4 false homozygotes and one false heterozygote) that was correctly assessed in repeated Sanger-sequencing using an alternative primer pair. In five additional variants, there was a zygosity status error in a single NGS experiment, which in 4/5 of cases was associated with low coverage (<20x).

In the clinical setting, false negatives are more concerning than false positives. Whereas false positives can be re-evaluated, detecting false negatives requires complete re-testing, which is unlikely to be routinely performed. The number of false negatives indicated are for ~1.3% of the exome, so over an entire exome the absolute number of false negatives will be correspondingly higher. We also note that these results overestimate the performance of NGS over the entire exome, since we purposely did not include genes in which the NGS is expected *a priori* to be less effective, e.g. genes that are GC-rich or known to have pseudogenes.

In summary, our results confirm the notion that neither Sanger-sequencing nor NGS can be regarded as a “gold-standard” method, as both had false positive and false negative calls. We have demonstrated that the performances of both NGS and Sanger-sequencing are satisfactory, and that no statistically significant differences were noted between the two sequencing methods with respect to the ability to sequence the targeted regions, and the observed, high (>98%) variant detection rates. Within NGS, it is evident that while clear differences were noted in the obtained sequencing coverage for the different capture kits, the detection rates of the true calls were not significantly different, at least when reaching the preset threshold of 70x coverage.

Bearing in mind the potential caveats that might be associated with pseudogenes or GC-rich regions, our data cautiously suggests that off-the-shelf exome-sequencing solutions might serve as a viable and practical alternative to gene(s) or gene-panel sequencing, albeit with a false negative rate of at least 1-3%. Nevertheless, physicians must be aware of the limitations of both Sanger-sequencing and NGS. As demonstrated herein, Sanger-sequencing primer binding-sites polymorphisms and chance or systematic NGS-coverage failure are integral hurdles of these methods. We show that Sanger-sequencing and different NGS solutions are synergistic, as any combination of the Sanger-sequencing, SureSelect and Nextera experiments yielded an overall greater detection rate. Accordingly, high index of suspicious for a given clinical diagnosis must overcome negative molecular results of either Sanger-sequencing or NGS and repeated investigation with an alternative method should be considered.

## ACKNOWLEDGMENTS

We thank the donor of the control sample.

## DECLARATIONS

### Availability of data and material

All data is available upon request

### Code availability

Not applicable

### Authors’ contributions

H.F., D.M.B. and E.L.L. conceived and designed the study. C.B., A.B., B.M., and L.F.S. provided the DNA samples to the study, managed and completed the genotyping campaign. M.E., Y.P., and D.A. completed the NGS analysis. H.F. and D.M.B wrote the paper. All authors discussed the results and commented on the manuscript.

