## Supplemental table 3 for "Performance comparison: exome-sequencing as a single test replacing Sanger-sequencing"

| **Supplemental Table 3.** Discordant variants**:** occurrence and analysis*^a^* | | | | | | | | | | | | | | | | |
| --- | --- | --- | --- | --- | --- | --- | --- | --- | --- | --- | --- | --- | --- | --- | --- | --- |
| **Gene** | **cDNA** | **rs number** | **Sanger*^b^*** | **SureSelect*^b^*** | | | | | **Nextera*^b^*** | | | | | | **True variant** | **Conclusion** |
|  |  |  |  | **Run 1** | | | | **Run 2** | **Run 1** | | | | **Run 2** | |  |  |
|  |  |  |  | **Buccal**  **dup A** | **Buccal**  **dup B** | **Blood dup A** | **Blood dup B** | **Blood** | **Buccal**  **dup A** | **Buccal dup B** | **Blood**  **dup A** | **Blood dup B** | | **Blood** |  |  |
| *ABCC6* | c.841A>G | rs4780606 | + | - | - | - | - | - | - | - | - | - | | - | N | Sanger FP |
| *ABCC6* | c.855C>T | rs4780605 | + | - | - | - | - | - | - | - | - | - | | - | N | Sanger FP |
| *ABCC6* | c.955A>G | rs72657699 | + | - | - | - | - | - | - | - | - | - | | - | N | Sanger FP |
| *IQCB1* | c.1301G>A | rs17849995 | - | + | + | + | + | + | + | + | + | + | | + | Y | Sanger FN |
| *OPA1* | c.473G>A | rs7624750 | - | + | + | + | + | + | + | + | + | + | | + | Y | Sanger FN |
| *TSC2* | c.5202T>C | rs1748 | - | + | + | + | + | + | + | + | + | + | | + | Y | Sanger FN |
| *TSC2* | c.5397G>C | rs1051771 | - | + | + | + | + | + | + | + | + | + | | + | Y | Sanger FN |
| *CFTR* | c.1251C>A | rs4727853 | - | - | + | - | + | + | - | + | + | + | | + | N | NGS FP |
| *NOTCH3* | c.2742A>G | rs1043997 | - | + | + | + | + | + | + | + | + | + | | + | N | NGS FP |
| *CACNA1A* | c.6669T>C | rs16051 | + | + | + | + | + | + | -0 | -0 | -0 | -0 | | -0 | Y | Nextera FN |
| *DLL3* | c.515T>G | rs8107127 | + | + | + | -0 | -0 | + | -0 | -0 | -0 | -0 | | -0 | Y | NGS FN |
| *DLL3* | c.546C>G | rs8106337 | + | + | + | -0 | + | + | -0 | + | -0 | -0 | | -0 | Y | NGS FN |
| *PIK3R2* | c.700A>C | rs2241088 | + | -0 | -0 | -0 | -0 | + | + | + | + | + | | -0 | Y | NGS FN |
| *COL5A1* | c.4560C>T | rs2228559 | + | + | + | -0 | + | + | + | + | + | + | | + | Y | SureSelect bl-R1DA FN |
| *COL6A2* | c.1333-8T>C | rs73159701 | + | -0 | -0 | + | + | + | + | + | + | + | | + | Y | SureSelect bu FN |
| *CDH1* | c.48+6C>T | rs3743674 | + | + | + | + | + | + | + | -0 | + | -0 | | -0 | Y | Nextera FN |
| *FRMD7* | c.1101T>C | rs7051368 | + | + | + | + | + | + | -0 | + | -0 | -0 | | + | Y | Nextera FN |
| *MEN1* | c.435C>T | rs61736636 | + | + | + | + | + | + | + | + | -0 | -0 | | + | Y | Nextera bl FN |
| *FGF3* | c.69G>T | rs41538178 | + | + | + | + | + | + | + | + | + | -0 | | + | Y | Nextera bl-R1DA FN |
| *GLIS3* | c.1270T>C | rs806052 | + | + | + | + | + | + | + | + | + | -0 | | + | Y | Nextera bl-R1DB FN |
| *COL6A2* | c.2979C>T | rs6652 | + | + | + | + | + | + | + | + | + | -0 | | + | Y | Nextera bl-R1DB FN |
| *LMX1B* | c.326+7G>C | rs1336980 | + | + | + | + | + | + | + | + | + | + | | - | Y | Nextera R2 FN |
| *NF1* | c.702G>A | rs1801052 | + | + | + | + | + | + | + | -0 | + | + | | + | Y | Nextera bu-R1DB FN |
| *COL5A2* | c.315C>A | rs4128539 | + | + | + | + | + | + | + | -0 | + | + | | + | Y | Nextera bu-R1DB FN |
| *EIF2AK3* | c.407C>G | rs867529 | + | + | + | + | + | + | - | + | + | + | | + | Y | Nextera bu-R1DA FN |
| *MESP2* | c.531G>A | rs75049807 | + | + | + | + | + | + | + | -0 | + | + | | + | Y | Nextera bu-R1DB FN |
| *MYH7* | c.1095G>A | rs735711 | + | + | + | + | + | + | - | + | + | + | | + | Y | Nextera bu-R1DA FN |
| *USH2A* | c.3812-8T>G | rs646094 | + | + | + | + | + | + | -0 | + | + | + | | + | Y | Nextera bu-R1DA FN |
| *GPC3* | c.338-5delT | rs370737647 | - | +0 | - | - | - | - | - | - | - | - | | - | N | SureSelect bu-R1DA FP |
| *MESP2* | c.558G>A | rs28546919 | - | - | - | - | - | +0 | - | - | - | - | | - | N | SureSelect R2 FP |
| *CFTR* | c.1365G>T | rs79074685 | - | - | - | - | - | - | - | - | +0 | - | | - | N | Nextera bl-R1DA FP |
| *COL1A1* | c.3046-6_ 3046-5delCT | rs138425306 | - | - | - | - | - | - | - | - | +0 | - | | - | N | Nextera bl-R1DA FP |
| *PIGN* | c.2620-5 delT | rs11437076 | - | - | - | - | - | - | - | - | +0 | - | | - | N | Nextera bl-R1DA FP |
| *APP* | c.10G>T | … | - | - | - | - | - | - | - | - | - | - | | +0 | N | Nextera R2 FP |
| *FRMD7* | c.575A>C | rs746689690 | - | - | - | - | - | - | - | - | - | - | | +0 | N | Nextera R2 FP |
| *NOTCH3* | c.4215C>T | … | - | - | - | - | - | - | - | - | - | - | | +0 | N | Nextera R2 FP |
| *NOTCH3* | c.4208G>T | … | - | - | - | - | - | - | - | - | - | - | | +0 | N | Nextera R2 FP |
| *SCNN1B* | c.275T>C | rs1596860414 | - | - | - | - | - | - | - | - | - | - | | +0 | N | Nextera R2 FP |
| *RAG1* | c.461delT | … | - | - | - | - | - | - | +0 | - | - | - | | - | N | Nextera bu-R1DA FP |
| *FOXC1* | c.1359_1361 delCGG | rs398123612 | - | - | - | - | - | - | - | +0 | - | - | | - | N | Nextera bu-R1DB FP |
| *EFNB1* | c.*731_*732 insC | rs61387446 | - | - | - | - | - | - | - | +0 | - | - | | - | N | Nextera bu-R1DB FP |
| *EFNB1* | c.*732_*733 insCAC | rs58007755 | - | - | - | - | - | - | - | +0 | - | - | | - | N | Nextera bu-R1DB FP |

*^a^*Shaded: Variants discordant between the Sanger and NGS experiments (N=13). Not Shaded: Variants discordant within the various NGS experiments (N=29).

*^b^* -: undetected variant, coverage >20x; -0: undetected variant, coverage <20x; +: detected variant, coverage >20x; +0: detected variant, coverage <20x or
Quality <100 or Quality/depth <7.

bl-blood; bu-buccal; D-duplicate; FP-False positive; FN-False negative; R-run; N-no; Y-yes.
